# Cdk1-mediated threonine phosphorylation of Sam68 modulates its RNA binding, alternative splicing activity, and cellular functions

**DOI:** 10.1101/2022.03.23.485498

**Authors:** Idir Malki, Inara Liepina, Nora Kogelnik, Adam Lightfoot, Oksana Gonchar, Hollie Watmuff, Andrew Bottrill, Andrew M. Fry, Cyril Dominguez

## Abstract

Sam68 is a member of the STAR family of proteins that directly link signal transduction with post-transcriptional gene regulation. Sam68 controls the alternative splicing of many oncogenic proteins and its role is modulated by post-translational modifications, including serine/threonine phosphorylation, that differ at various stages of the cell cycle. However, the molecular basis and mechanisms of these modulations remain largely unknown. Here, we combined mass spectrometry, NMR spectroscopy, and cell biology techniques to provide a comprehensive post-translational modification (PTM) mapping of Sam68 at different stages of the cell cycle in HEK293 and HCT116 cells. We established that Sam68 is specifically phosphorylated at T33 and T317 by Cdk1, and demonstrated that these phosphorylation events reduce the binding of Sam68 to RNA, control its cellular localization, and reduce its alternative splicing activity, leading to a reduction in the induction of apoptosis and an increase in the proliferation of HCT116 cells.

## INTRODUCTION

Alternative splicing of pre-mRNA is a highly regulated mechanism that allows for the generation of proteomic diversity from a limited number of human genes (1). Cell signaling plays a major role in the regulation of alternative splicing (2,3) and splicing factor mutations or copy number variations are associated with many genetic diseases, including cancer (4,5).

A typical example is the splicing factor Sam68 (Src-associated protein during mitosis of 68kDa) that controls the alternative splicing of many pre-mRNAs encoding for oncogenic proteins, such as cyclin D1, CD44, and Bcl-x (6-8). Sam68 plays important roles in cell cycle regulation, apoptosis and viral replication, through its functions in various aspects of gene expression, including transcription, pre-mRNA splicing and RNA export (9,10). Sam68 is also an oncogene (10,11): high expression of Sam68 correlates with poor prognosis in many cancers, including prostate, and colon cancers (12,13), while depletion of Sam68 decreases cell migration in HeLa cells (14), and inhibits proliferation and tumor progression in prostate and breast cancers (12,15). Importantly, Sam68’s cellular functions are regulated by cell signaling through post-translational modifications (PTMs), providing a direct link between cell signaling and post-transcriptional gene regulation (9,10).

Sam 68 belongs to the STAR (Signal Transduction and Activation of RNA) family of proteins (16,17) and contains a central QUA1/KH domain that is responsible for its homodimerization and RNA binding. This domain is flanked by N- and C-terminal regulatory regions, such as a tyrosine-rich region (YY) targeted by tyrosine kinases, proline-rich regions (P) responsible for binding SH3 and WW domain-containing proteins, Arg/Gly (RG)-rich regions targeted by arginine methyltransferases and a nuclear localization signal (NLS). We have recently revealed the structural basis of RNA recognition and dimerization by the QUA1/KH domain and proposed a model for the role of Sam68 in alternative splicing whereby Sam68 dimers loop out regions of its target pre-mRNAs to induce either exon inclusion or skipping (18).

It is well known that the cellular functions of Sam68 are controlled by PTMs, including tyrosine, serine and threonine phosphorylation by the Src kinase family (19,20), Erk1 (7), and Cdk1 (21), arginine methylation by PRMT1 (22), sumoylation by PIAS1 (23), lysine acetylation by CBP (24) and deacetylation by HDAC6 (25). For example, phosphorylation of Src-like kinases enhances the production of the oncogenic isoform of Bcl-X (8).

While tyrosine phosphorylation and arginine methylation of Sam68 have been well documented, serine/threonine (S/T) phosphorylation of Sam68 remains largely unexplored (26). Previous research in HeLa and NIH3T3 cells showed that serine residues of Sam68 are phosphorylated throughout the cell cycle but that threonine phosphorylation occurs only during mitosis and is mediated by CDK1 (21). Mutation of a putative Cdk1 target site (T317) reduced the levels of murine Sam68 phosphorylation by Cdk1 in vitro by 30%, suggesting that this is likely to be one site of phosphorylation but that at least one other phosphorylation site is present in Sam68. Precisely which residues of Sam68 are phosphorylated by Cdk1 and the functional consequences of these phosphorylation events remain unknown. Later, Sam68 was shown to be phosphorylated by Erk1 in EL4 and SW480 cells and that this phosphorylation event reduced Sam68’s RNA binding ability (27) but stimulated the alternative splicing activity of Sam68 towards CD44 exon 5 inclusion (7), the splicing of the *Cyclin D1b* isoform (6) and SRSF1 nonsense mRNA decay (28). A triple mutant S58A/T71A/T84A reduced the stimulation of CD44 exon v5 inclusion while single mutants did not, suggesting that the phosphorylation of at least one of these three residues is important for splicing stimulation (7). However, there is no direct evidence that these three residues are phosphorylated by Erk1. Furthermore, this study was done on murine Sam68 and T71 is not conserved in human Sam68. Finally, Sam68 was shown to be phosphorylated at S20 following depolarization of neuronal cells, possibly by CAMK IV, leading to alterations in the splicing of Neurexin 1 (29), but it remains unclear whether this phosphorylation event is restricted to neuronal cells or is also present in other cell types.

Here we used mass spectrometry to map the serine and threonine residues of Sam68 phosphorylated at different stages of the cell cycle in HEK293 and HCT116 cells. In addition, we have characterized the specific phosphorylation of Sam68 at T33 and T317 by Cdk1 showing that their phosphorylation not only reduce Sam68’s ability to bind RNA and regulate splicing, but also modulate its cellular localization and functions in apoptosis and proliferation. We propose a model whereby the intrinsically disordered regulatory regions of Sam68 anchor the protein to its target pre-mRNAs, thus increasing its affinity for RNA. However, during mitosis, phosphorylation of Sam68 by Cdk1 reduces its affinity for RNA and allows for competing splicing factors to bind the pre-mRNA, resulting in switches in RNA splicing patterns.

## RESULTS

### Mapping of Sam68 PTMs in HEK293 and HCT116 cells

The amino acid sequence of Sam68 contains 28 serines and 19 threonines that are mainly located in the N-terminal and C-terminal regulatory regions (Figure 1A). To map serines and threonines phosphorylated in cells, flag-tagged full-length human Sam68 was transfected in HEK293 and HCT116 cells. 48 hours post-transfection, asynchronous cells were harvested and Flag-Sam68 was purified using a flag antibody, and digested with trypsin. Post-translational modifications (PTMs) were identified by LC-MS/MS (Table 1). The datasets did not provide any sequence coverage of the tyrosine-rich region of Sam68, and no phosphotyrosine residues were detected. This was expected, as digestion with trypsin gives very large peptides for this region. However, as expected, we observed methylation of multiple arginines and acetylation of multiple lysines as previously reported (22,24). In terms of serine and threonine (S/T) phosphorylation, we observed phosphorylation of either S18 or S20 (S18/S20), either T33 or S35 (T33/S35) in HEK293T cells and either S18 or S20, either T33 or S35 and S113 in HCT116 cells (Table 1).

**Table 1:** List of Sam68 post-translational modifications identified by LC-MS/MS.

|  | HCT116<br>unsynchronized | Hek293<br>unsynchronized | HEK293<br>S-phase | HEK293<br>M-phase | HEK293<br>G1-phase |
| --- | --- | --- | --- | --- | --- |
| Arginine<br>methylation | R43, R45, R52,<br>R282, R284,<br>R289, R291,<br>R302, R304,<br>R310, R315,<br>R320, R325,<br>R331, R340 | R36, R186,<br>R282, R284,<br>R289, R291,<br>R302, R304,<br>R310, R325,<br>R331, R340 | R4, R43, R45, R52,<br>R186, R213, R282,<br>R284, R289, R291,<br>R302, R304, R310,<br>R320, R325, R331,<br>R340, R345 | R43, R45, R52,<br>R186, R282, R284,<br>R289, R291, R302,<br>R304, R310, R315,<br>R320, R325, R331,<br>R340, R345 | R282, R284,<br>R289, R291,<br>R302, R325,<br>R331 |
| Lysine Acetylation | K111, K139 | K139, K194,<br>K206 | K131, K139, K208 | K111, K131, K138,<br>K139, K152, K206,<br>K208, K432 | K111, K131,<br>K139, K169,<br>K175, K194,<br>K206, |
| Serine/Threonine<br>Phosphorylation | <u>S18/S20*</u>,<br>S35/T33**, S113 | S18/S20*,<br>S35/T33** | S20 | S18/S20*, T33,<br>S150, T317 | S20, S58, S70 |

\* We observed single phosphorylation of the peptide containing S18 and S20, indicating that we can not differentiate which serine is phosphorylated (HCT116), or that two populations co-exist in the same sample one with S18 phosphorylated, another with S20 phosphorylated (HEK293 unsynchronized and M-phase).
\*\* We observed single phosphorylation of the peptide containing S35 and T33, indicating that we can not differentiate which amino acid is phosphorylated (HCT116), or that two populations co-exist in the same sample one with S35 phosphorylated, another with T33 phosphorylated (HEK293).

**Figure 1:**
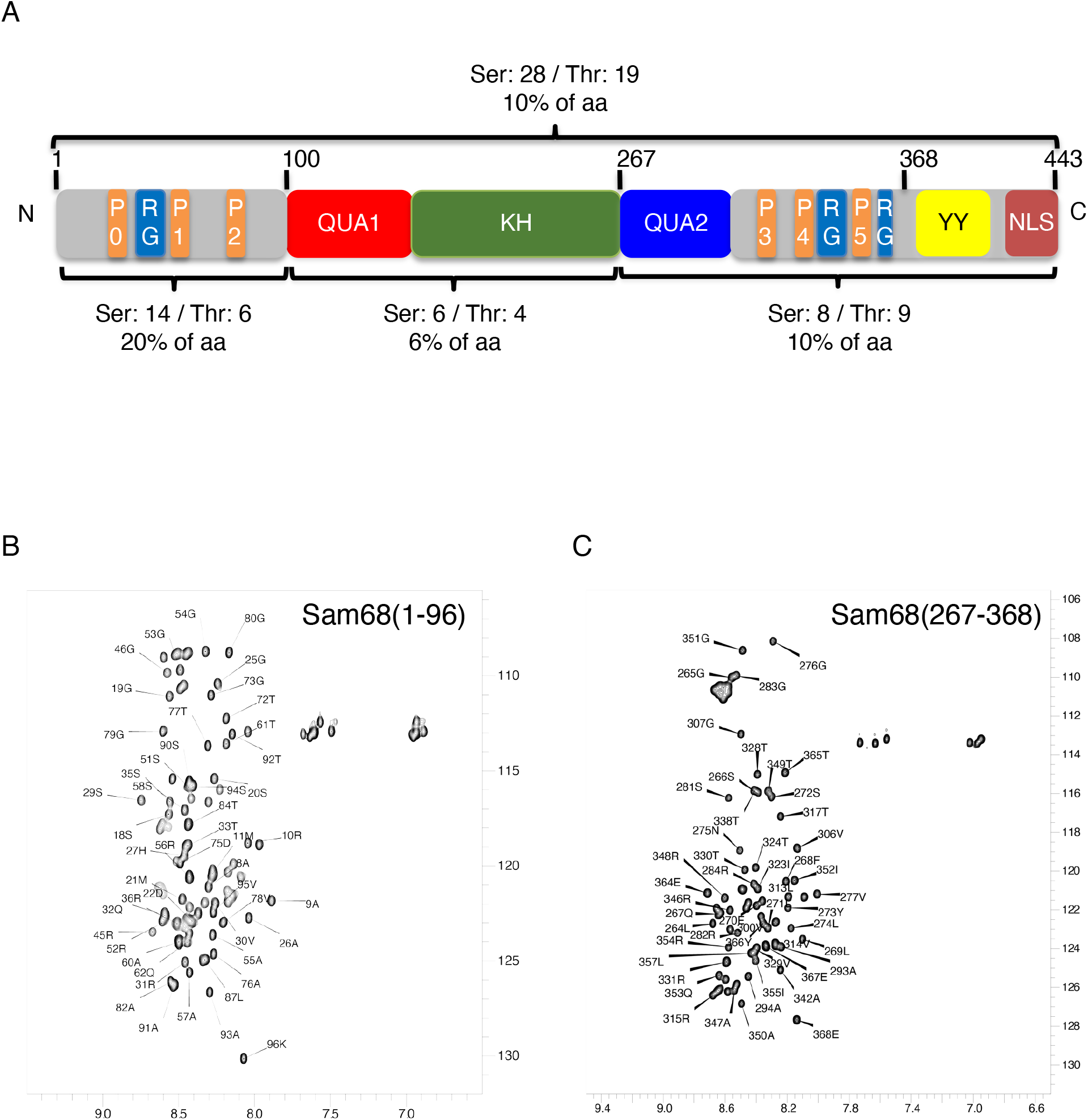
Domain organization of Sam68 and NMR analysis of the N-terminal and C-terminal regions. (**A**) Sam68 is a 443-amino acids protein. The QUA1 and KH domains are responsible for dimerization and RNA binding. The N-terminal region (residues 1-96), the QUA2 region (267-283) and the C-terminal region (residues 284-443) are predicted to be intrinsically disordered and contain regulatory motifs such as Pro-rich motifs (P0 to P5), RG-rich motifs (RG), a tyrosine-rich region (YY) and a nuclear localization sequence (NLS). The number and percentage of serine and threonine for each region/domain is indicated. (**B, C**) NMR (^1^H-^15^N)-HSQC spectra of Sam68 N-term (residues 1-96) (**B**) and C-term (residues 267-368) at 4°C (**C**). The assignment of the backbone amide resonances is indicated.

Next, we investigated the PTMs of Sam68 in HEK293 cells at different stages of the cell cycle by treating cells with either nocodazole (G1 and M phases) or with hydroxyurea (S phase). The enrichment in specific phases of the cell cycle was verified by flow cytometry (Supplementary Figure S2) and samples were analysed by LC-MS/MS. S18/S20 phosphorylation was observed in all stages of the cell cycle. Additionally, S58 and S70 were phosphorylated during G1. These two specific phosphorylation events could be mediated by Erk1 as previously suggested (7). Interestingly, we identified T33 and T317 phosphorylation in mitosis. These phosphorylation events could be mediated by Cdk1/Cyclin B (21).

### Cdk1 phosphorylates Sam68 at T33 and T317 in vitro

To precisely investigate the phosphorylation of Sam68 by Cdk1 *in vitro*, we used NMR spectroscopy that is very powerful for the study of PTMs at atomic resolution (30).

The N-terminal (residues 1-96; N-term) and C-terminal (residues 267-368; C-term) regions of Sam68 were expressed in *E. coli* in the presence of ^15^N-ammonium chloride and ^13^C-glucose before purification. The NMR spectra show that these regions are intrinsically disordered and 72% and 65% of the non-proline residue backbone resonances for the N-term and the C-term regions, respectively, could be unambiguously assigned (Figure 1B, C). Next, these purified regions were incubated with ATP, MgCl_2_, and active Cdk1/cyclin B. Phosphorylation was quantitatively monitored by recording a (^1^H-^15^N)-HSQC at time zero at 4°C, followed by consecutive 2D (^1^H-^15^N)-SOFAST-HMQC NMR experiments for 16 hours at 20°C, and a final (^1^H-^15^N)-HSQC at 4°C. During the experiment, the resonance peaks of T33 and T317 shifted downfield in both ^1^H and ^15^N dimension, a typical signature of serine or threonine phosphorylation (31) (Figure 2A, B). Resonances corresponding to other threonine and serine residues were not affected by the addition of Cdk1/cyclin B, indicating that only T33 and T317 were phosphorylated by Cdk1 *in vitro*. Incubation of the N-term with Cdk1 induced chemical shift perturbations of residues surrounding T33 (notably H27, S29, R31, Q32, R35 and Q36) (Figure 2C), suggesting a possible conformational change upon T33 phosphorylation. In contrast, phosphorylation of T317 in the C-term region only induced minor chemical shift changes to the surrounding residues, suggesting no major conformational changes (Figure 2D).

**Figure 2:**
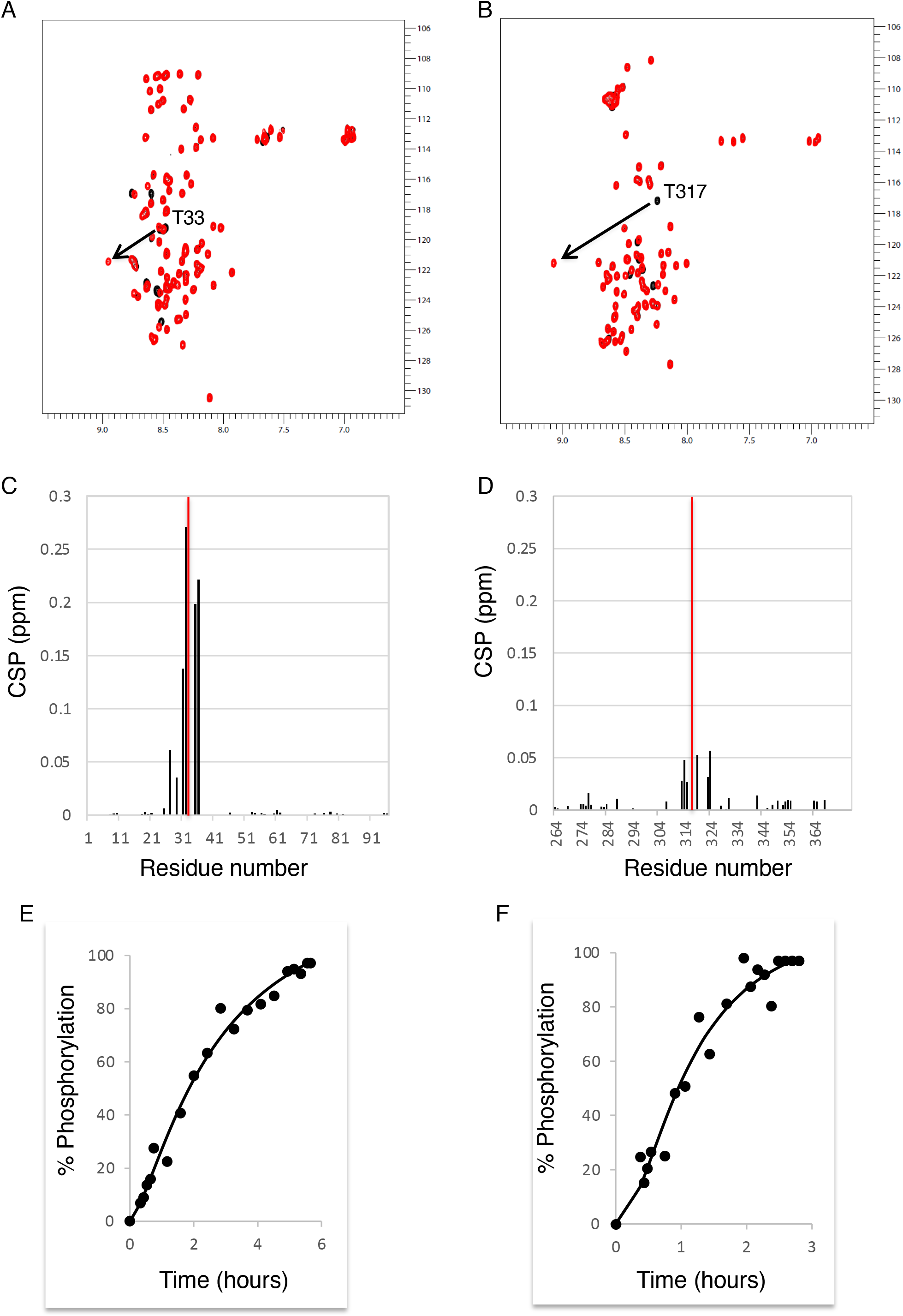
Sam68 N-term and C-term are phosphorylated at T33 and T317 by Cdk1. (**A, B**) (^1^H-^15^N)-HSQC spectra overlay of Sam68 N-term (**A**) and C-term (**B**) before (black) and after 16 hours incubation with commercial active Cdk-1/Cyclin B (red) at 4°C. T33 and T317 resonance peaks are indicated. (**C, D**) Chemical shift perturbation of backbone amides as a function of the N-term (**C**) and C-term (**D**) amino acid sequence upon Cdk1 phosphorylation. For clarity, the CSP of T33 and T317 are represented as red bars and are not to scale. (**E, F**) Normalized intensity of phosphorylated T33 (**E**) and T317 (**F**) backbone amide peak as a function of time after Cdk1/cyclin B addition. The data were fitted using a Hill function.

Analysis of the (^1^H-^15^N)-SOFAST-HMQC spectra at different times after addition of Cdk1 allowed us to investigate the kinetics of T33 and T317 phosphorylation by Cdk1 (Figure 2E, F). Interestingly, phosphorylation of these residues did not follow a Michaelian response but a sigmoidal ultrasensitive response. Such a response is not uncommon in cell signaling systems (32,33) and has already been suggested for other Cdk1 substrates (34-36). Overall, the complete phosphorylation of T317 is faster (2.5 hours) than T33 (5 hours). This is consistent with the amino acid sequence surrounding the phosphorylated residues, T_33_PSR and T_317_PVR. The target sequence for Cdk1 is (S/T)PX(K/R), where X can be any amino acid. However preferential amino acids have been described at this position with a valine predicted to provide a better Cdk1 target than a serine (37).

### The N-terminal and C-terminal regions of Sam68 bind RNA

Like other STAR proteins, Sam68 binds predominantly and specifically to RNA through its central STAR domain (18,38-40). However, the dissociation constant of full-length Sam68 to SELEX-derived RNAs was reported to be in the low nanomolar range (41,42), while we found that the isolated STAR domain binds the same RNA sequences with dissociation constants in the low micromolar range (18). This suggests that regions of Sam68 outside the STAR domain could contribute to RNA binding. Sam68 N-terminal and C-terminal regions contain multiple RG-rich regions and it is well established that RG-rich regions of many proteins have RNA binding properties (43). Accordingly, it has been shown that Sam68’s RG-rich regions are capable of binding RNA non-specifically (44).

We therefore used NMR spectroscopy to investigate the binding of purified Sam68 N-term and C-term proteins with the RNA sequence G8-5 that was previously identified as a high affinity Sam68 binder by SELEX (41) (Figure 3A). Increasing the G8.5 molar ratio with Sam68 N-term and C-term induced chemical shift perturbations consistent with the formation of a complex between these regions and the RNA (Figure 3B, C). These results demonstrate that the Sam68 N-term and C-term regions can bind RNA independently of the rest of the protein. Analysis of the titration data suggests that both domains bind the G.8 RNA with dissociation constants in the low micromolar range (1-10μM for N-term and 30-70μM for C-term) (Supplementary Figure S3), very similar to the STAR domain affinity for the same RNA sequence (18).

**Figure 3:**
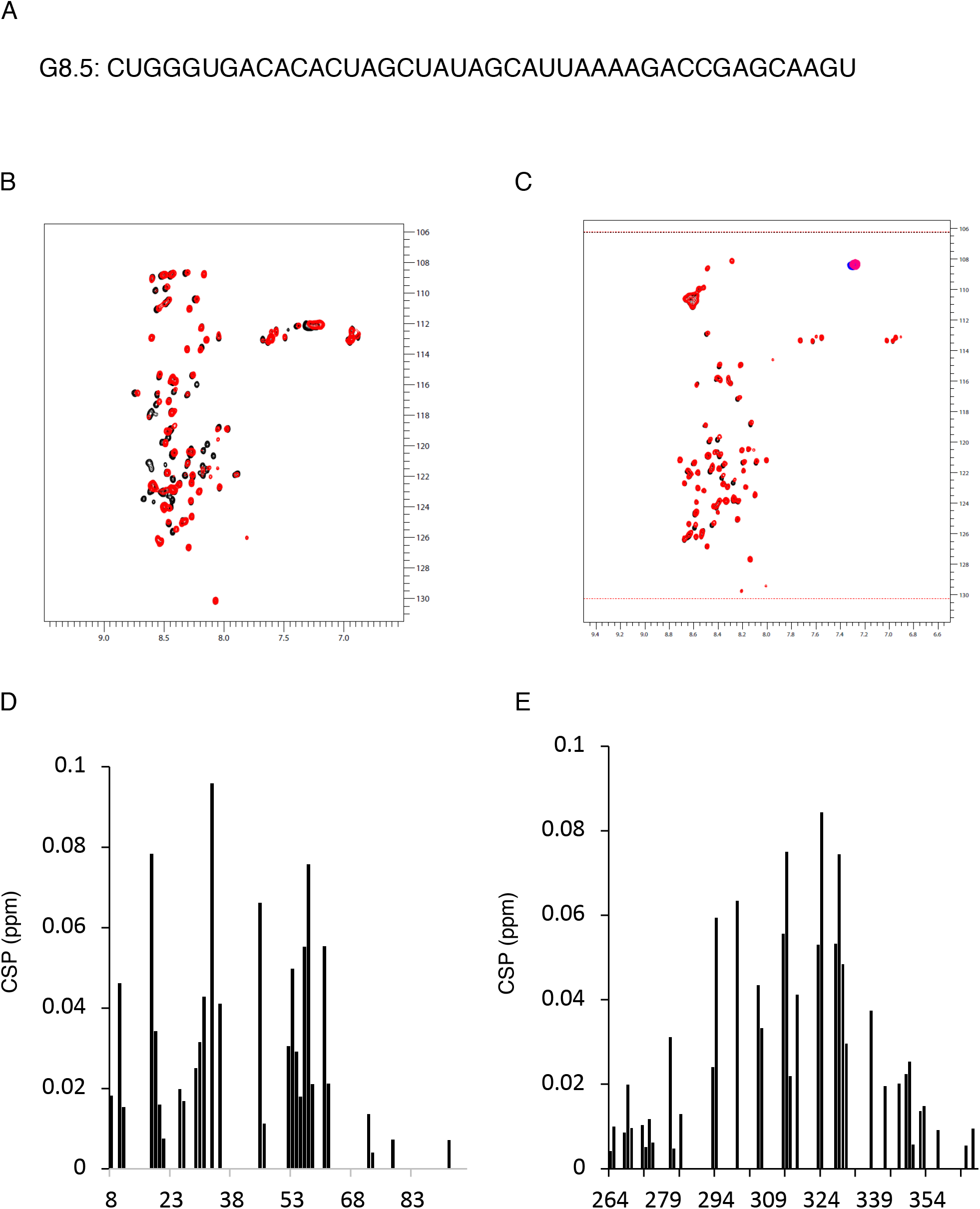
The N-term and C-term regions of Sam68 bind RNA. (**A**) nucleotide sequence of the G8.5 RNA identified previously as a high-affinity binder of Sam68 (41). (**B, C**) (^1^H-^15^N)-HSQC spectra of Sam68 N-term (**B**) and C-term (**C**) before (black) and after (red) addition of excess G8.5 RNA (protein:RNA molar ratio of 1:2) at 4°C. (**D, E**) Chemical shift perturbation of Sam68 N-term (**D**) and C-term (**E**) backbone resonances upon RNA interaction as a function of the amino acid sequence.

Analysis of the chemical shift mapping shows that RNA binding affects most residues of the N-term and C-term, but in particular residues 10-62 for the N-term and 281-342 for the C-term (Figure 3D, E). This is consistent with the presence of multiple RG-rich motifs in the primary sequence, suggesting non-specific binding of these RG-rich regions to the RNA. However, it could also indicate that RNA binding induces conformational changes to Sam68 N-term and C-term. Interestingly, the chemical shifts of T33 and T317 are both significantly perturbed by the addition of RNA demonstrating that their chemical environment is affected either by direct binding to the RNA or by RNA-induced conformational changes. Moreover, this implies that phosphorylation of these residues could modulate RNA-binding through the N-term and C-term regions.

### Phosphorylation of Sam68 N-terminal and C-terminal regions by Cdk1 decreases their RNA binding ability

We next investigated whether the phosphorylation of T33 and T317 by Cdk1 had an effect on the RNA binding ability of Sam68 N-term and C-term. For this purpose, we measured NMR (^1^H-^15^N)- HSQC spectra of Cdk1-phosphorylated Sam68 N-term and C-term regions in the absence or the presence of an excess G8.5 RNA (Figure 4A, B). The phosphorylated forms of Sam68 N-term and C-term were still able to bind the G8.5 RNA but the intensity of the chemical shift perturbations were strongly reduced compared to the non-phosphorylated regions (Figure 4C, D). This indicates that phosphorylation of T33 and T317 reduces but does not abolish RNA binding. Together, our data demonstrate that the N-terminal and C-terminal regions of Sam68 are capable of binding RNA and that phosphorylation of T33 and T317 by Cdk1 reduces their RNA binding ability.

**Figure 4:**
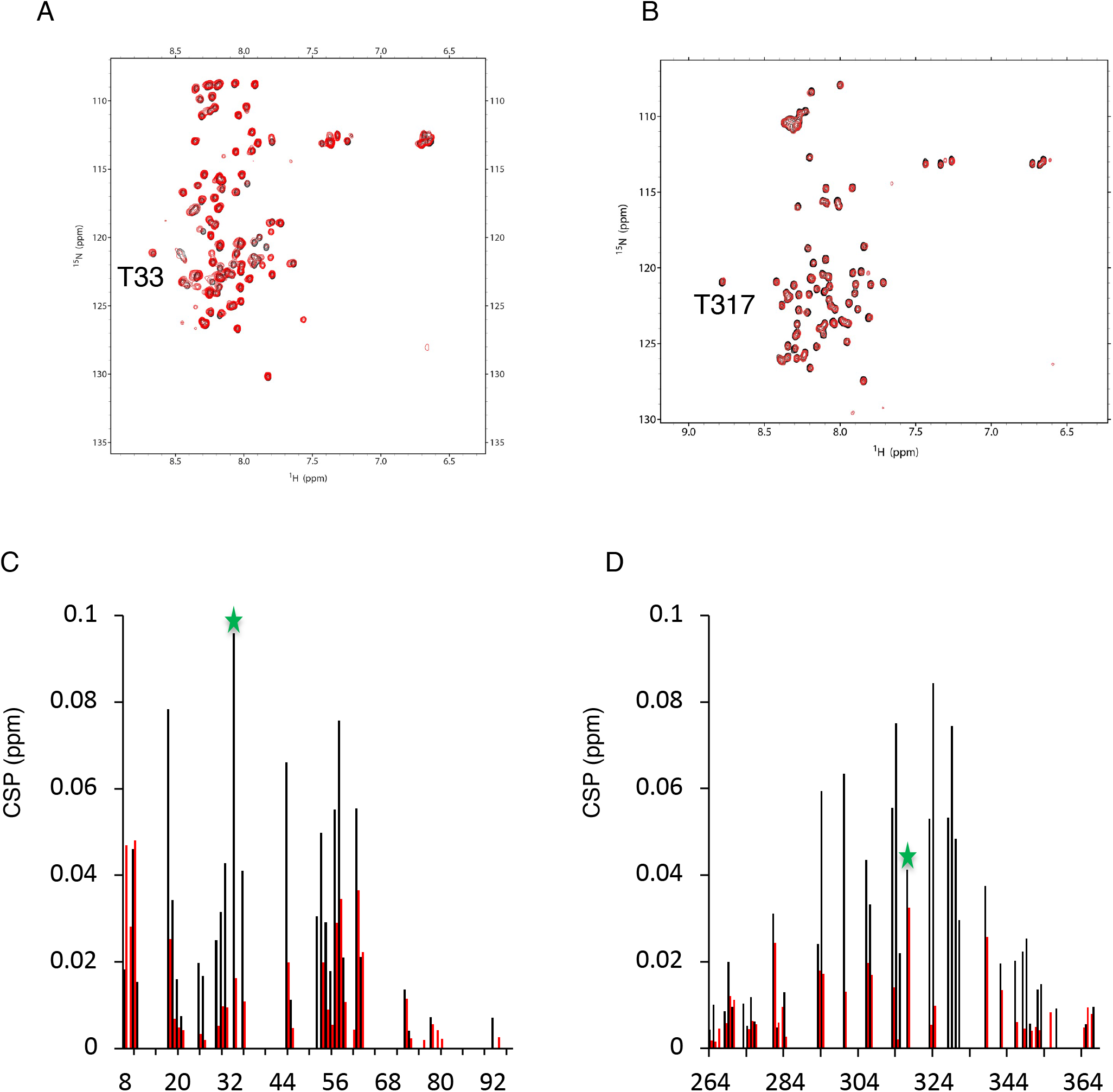
T33 and T317 phosphorylation reduce RNA binding ability of Sam68 N-term and C-term. (**A, B**) (^1^H-^15^N)-HSQC spectra of Cdk1-phosphorylated Sam68 N-term (**A**) and C-term (**B**) before (black) and after (red) addition of 2 molar equivalent of G8.5 RNA at 4°C. Resonances of T33 and T317 are indicated. (**C, D**) Chemical shift perturbation of Sam68 N-term (**C**) and C-term (**D**) in their unphosphorylated (black) or Cdk-1 phosphorylated (red) forms upon RNA binding. T33 and T317 CSPs are labeled with a green star.

### T33 and T317 phospho-mimetic mutations alter the cellular localization of Sam68

Sam68 is diffusely distributed throughout the nucleus in HEK293, MEF, HF-7650, REF-52 cells, while in some cell lines such as HeLa, BT-20, Hs578T, Sam68 is also localized in nuclear speckles, called Sam68 Nuclear Bodies (SNBs) (45). The percentage of cells displaying SNBs is above 85% in HeLa and BT-20, while it is only 4.5% in HEK293 and below 1% in MEF, HF-7650, REF-52 cells. Single amino acid mutation and deletion of some regions of Sam68 (notably RNA-binding mutants in the STAR domain) result in the relocalization of Sam68 to different compartments. This has led to the description of seven localization patterns for wild-type and mutant Sam68 ranging from diffuse in the nucleus with some SNBs to only SNB or even fibrous cytoplasmic structures. Sam68 mutants display varied percentages of each pattern (45). Notably, all mutations in the KH RNA binding domain of Sam68 led to a significant decrease in the number of cells displaying a diffuse nuclear localization and an increase in the number of cells displaying nuclear and/or cytoplasmic speckles, indicating that any interference with the RNA binding properties of Sam68 lead to increase SNB formation and/or cytoplasmic localization (45).

To investigate the consequences of Sam68 T33 and T317 phosphorylation on cellular localization, we created phospho-mimetic GFP-tagged Sam68 mutants (T33E, T317E, and T33E/T317E) and compared their localization in HEK293 and HCT116 (Figure 5A, B). No changes in localization of WT and mutant Sam68 was observed in HEK293. However, in HCT116, significant changes were observed (Figure 5C). As expected, WT Sam68 is mainly diffused in the nucleus with sometimes few SNBs (76.2±6.3%, pattern A), with a small proportion of cells displaying Sam68 exclusively in SNBs (9.7±5.7%, pattern B), or localized in both the nucleus and the cytoplasm (9.0±4.6%, pattern C). In contrast, the phospho-mimetic mutants displayed significant localization differences when compared to WT Sam68 (Figure 5C). These changes are more pronounced with the phospho-mimetic mutants T33E and T33E/T317E that exhibit a two-fold increase in the percentage of cells displaying pattern B (19.9±2.46 and 23.2±4.5). The double T33E/T317E mutant also exhibits a significant decrease in the percentage of cells with pattern A (60.1±4.9). These changes in localization are similar to previous mutations in the KH RNA binding domain of Sam68 (45). Taken together with our NMR results, this localization data supports the hypothesis that phosphorylation of Sam68 by Cdk1 reduces its affinity for RNA.

**Figure 5:**
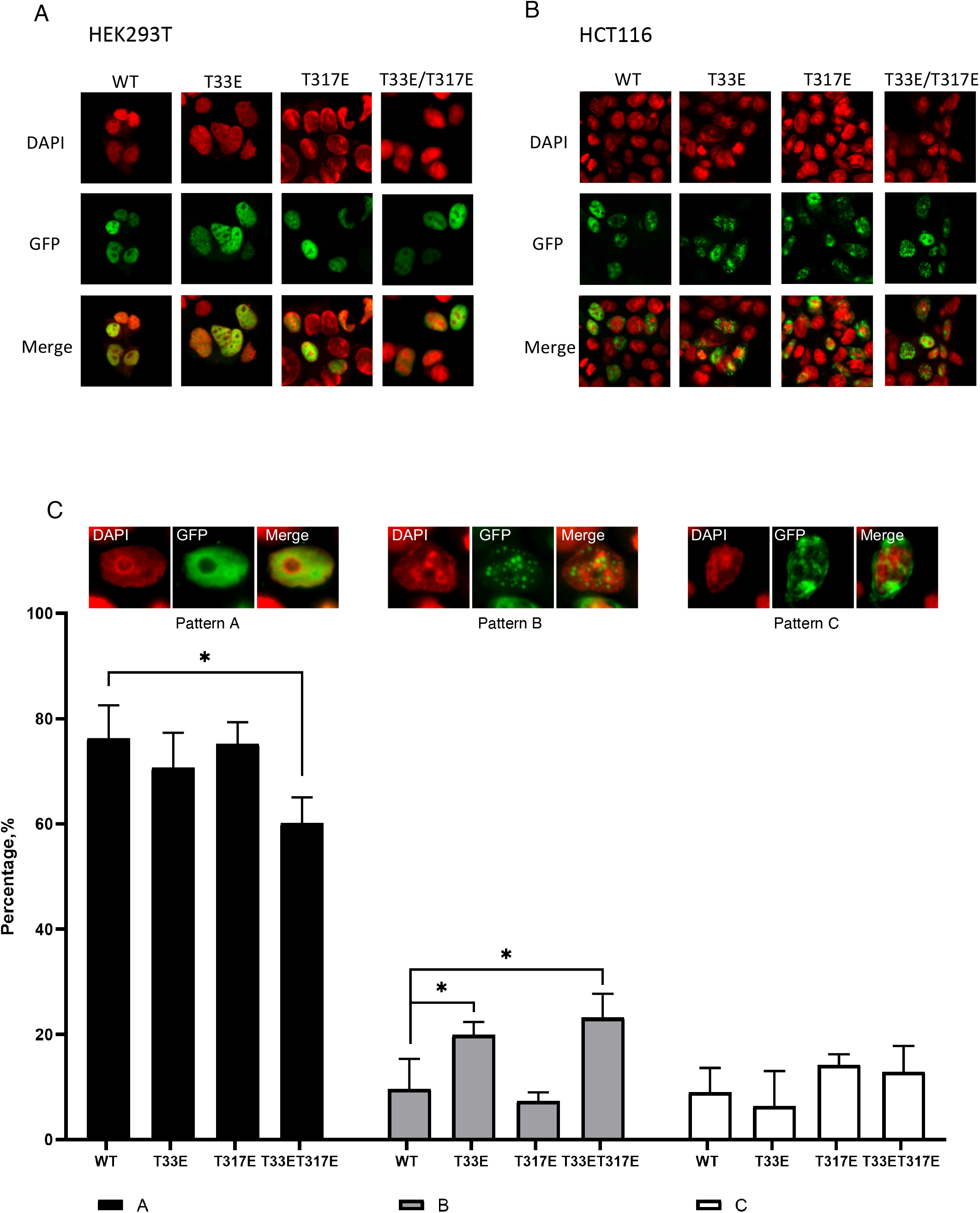
Cellular localization of Sam68 wild-type (WT), T33E, T317E and T33E/T317E mutants in HEK-293T and HCT-116 cells. (**A, B**) Confocal fluorescence images of HEK-293T (**A**) and HCT-116 (**B**) cells transfected with either GFP-tagged Sam68 WT, or phospho-mimetic mutants. DAPI is shown in red and GFP in green. (**C**) Representative images of 3 different classes (patterns A-C) of Sam68 wild-type and mutants cellular localization in HCT-116 cells (Top) and percentage of cells displaying Sam68 or mutants localized in each class for 50-100 cells analysed per sample (Bottom).

### T33 and T317 phospho-mimetic mutants have reduced Sam68 splicing activity

To test whether phosphorylation of Sam68 at T33 and T317 might affect its splicing activity, we investigated the ability of the phospho-mimetic mutants to regulate splicing of CD44 and Bcl-x minigenes (7,8) in HCT116 cells (Figure 6). As previously reported, transfection of wild-type Sam68 induced the inclusion of exon v5 in CD44 (Figure 6A) and increased the Bcl-x_S_/Bcl-x_L_ ratio (Figure 6B). However, the phospho-mimetic mutants were significantly less efficient at splicing CD44 exon v5 compared to WT Sam68 (0.41±0.05, 0.44±0.02 and 0.40±0.01 for T33E, T317E and T33E/T317E, respectively, compared to 0.52±0.03 for WT Sam68) and at shifting splicing towards the X_S_ isoform (X_S_/X_L_ ratio of 0.12±0.01 for T33E/T317E compared to 0.19±0.03 for WT Sam68). Hence, phosphorylation of Sam68 by Cdk1 modulates its regulatory activity on CD44 and Bcl-x splicing.

**Figure 6:**
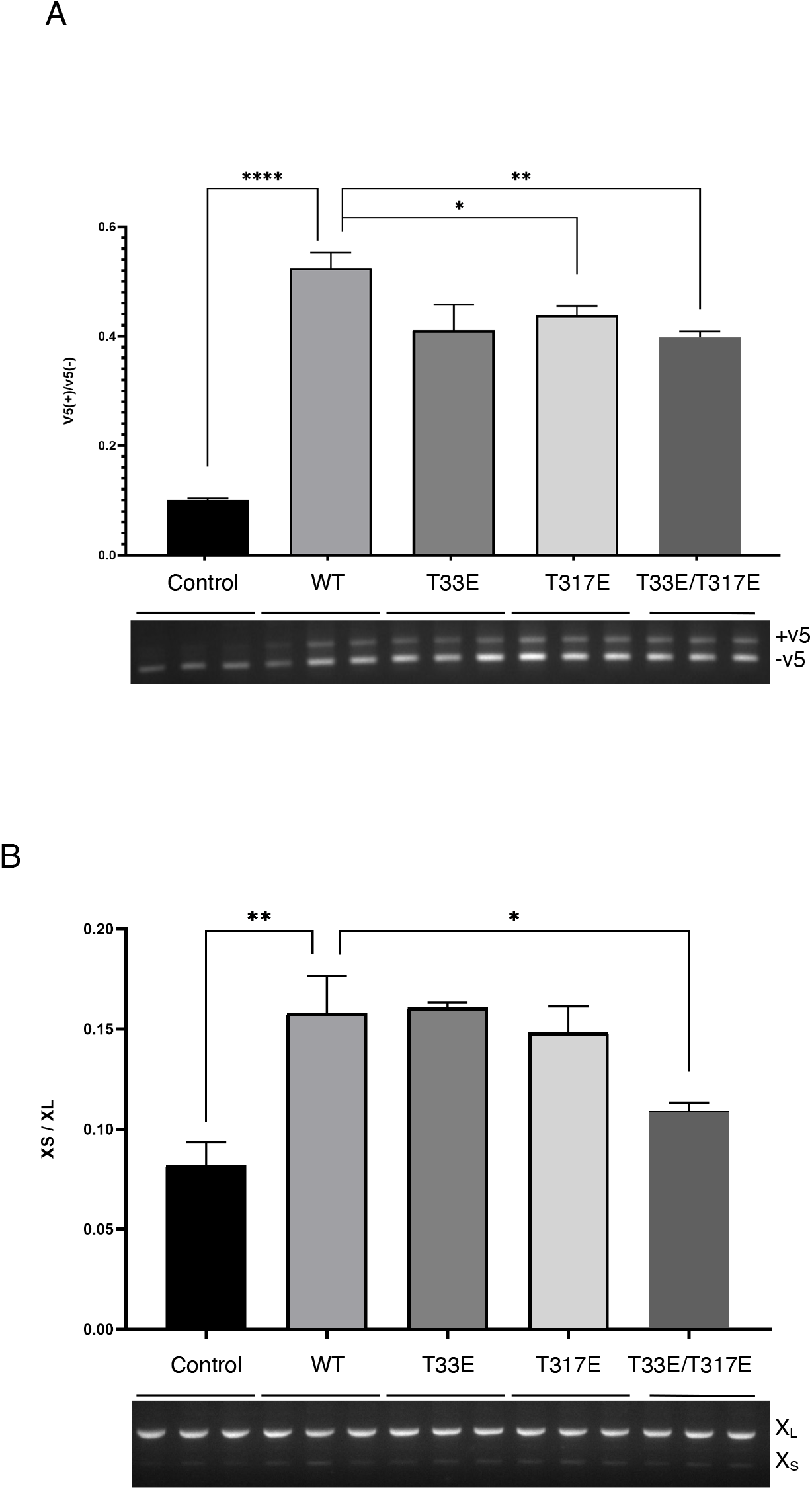
Splicing activity of Sam68 WT and phospho-mimetic mutants on CD44 exon v5 and Bcl-x minigenes in HCT-116 cells. Effect of Sam68 WT, T33E, T317E and T33E/T317E transfection on the alternative splicing of CD44 exon v5 (**A**), and Bcl-x (**B**) minigenes. Bottom: agarose gel electrophoresis showing splicing of the minigenes in response to co-transfected proteins. Top: quantification of biological replicates from three independent co-transfection experiments. Bar chart plotting and analysis were performed using GraphPad Prism. Error bars represent the standard deviation of three independent experiments. P values were calculated using an independent two-sample t-test (*: p<0.05, **: p<0.005 and ****: p<0.00005). Uncropped gels are shown in Supplementary Figure S8.

### T33 and T317 phospho-mimetic mutants affect cell cycle progression, decrease apoptosis and increase proliferation

Sam68 is a key regulator of cell cycle progression and apoptosis(46). We therefore performed flow cytometry with DNA staining to investigate the consequences of transfecting wild-type (WT) Sam68 and phospho-mimetic mutants on HCT-116 cell cycle progression (Supplementary Figure S4). Significant changes could be observed in the relative proportions of sub-G1, G1 and G2/M populations upon expression of either WT Sam68 or the phospho-mimetic mutants (Figure 7A, B). Expression of Sam68 WT significantly decreased (from 27.4±0.9% to 19.7±0.2%) the G2/M population and significantly increased (from 0.9±0.1% to 12.3±1.4%) the sub-G1 population compared to untransfected cells. In contrast, expression of Sam68 T33E, T317E and T33E/T317E mutants resulted in increased G2/M populations (26.6±0.8%, 22.6±0.9%, and 21.8±0.5%, respectively) compared to WT expression. Moreover, Sam68 T33E transfection resulted in a significant decrease in the sub-G1 population (5.1±0.6%) and an increase in the G1 population (36.5±2.1%) compared to WT.

**Figure 7:**
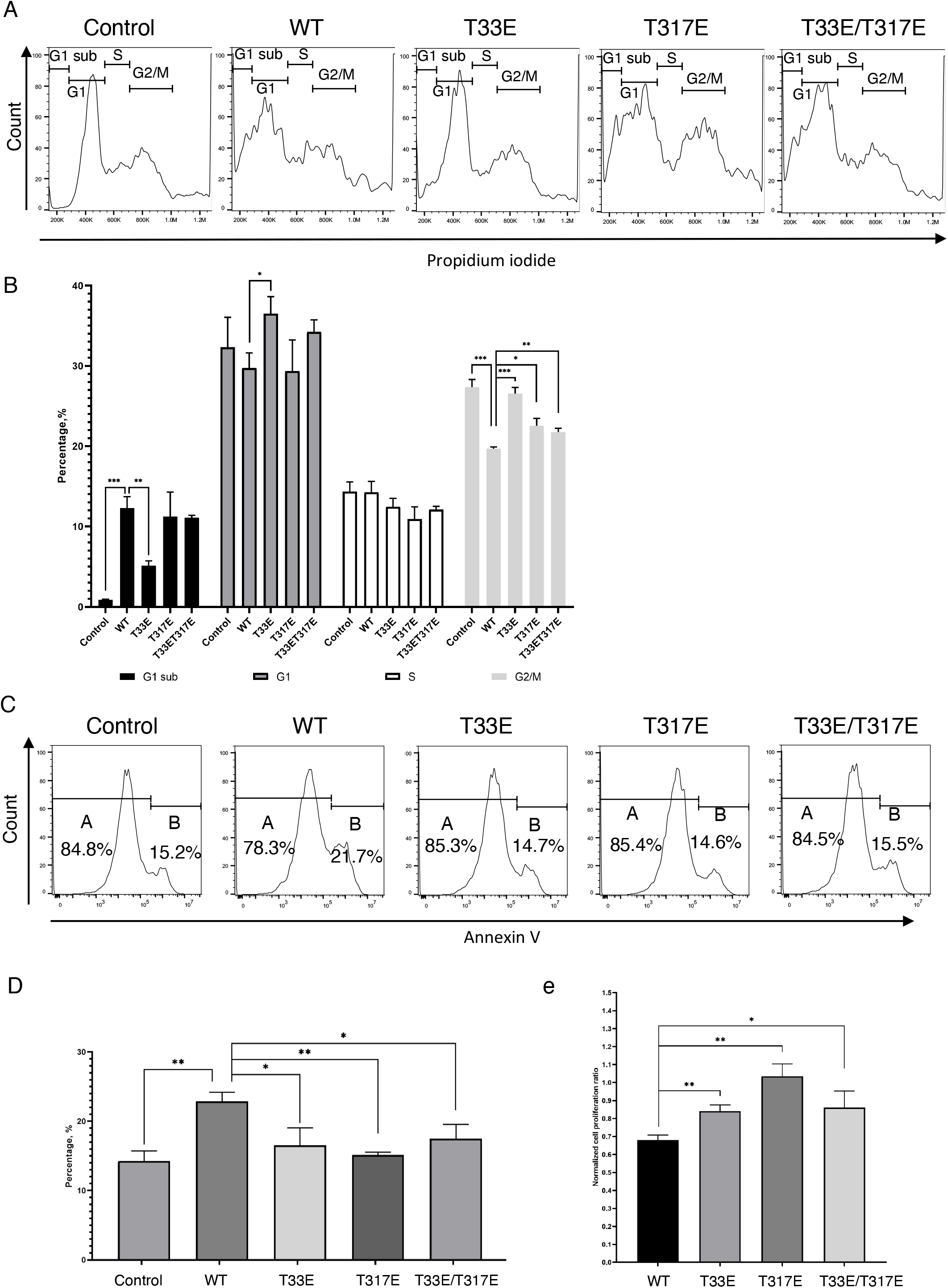
Effect of Sam68 WT or phospho-mimetic mutants on cell cycle progression, apoptosis and proliferation of HCT116 cells. (**A, B**) Effect of Sam68 WT and mutants transfection on cell cycle progression. (**A**) Flow cytometry images and (**B**) percentage of cells in sub G1 (black), G1/G0 (dark grey), S (white), and G2/M (light grey) phases 48 hours after transfection with Sam68 WT or mutants. (**C, D**) effect of Sam68 WT and mutants transfection on cell apoptosis. (**C**) Flow cytometry images and (**D**) percentage of apoptotic cells 48 hours after transfection with Sam68 WT or mutants. (**E**) Normalized cell proliferation increase between 48 and 72 hours after transfection with Sam68 WT or mutants. Bar chart represents the ratio between the number of GFP-positive cells at 72 hours over 48 hours. Error bars represent the standard deviation of three independent experiments. P values were calculated using an independent two-sample t-test (statistical significance shown as: *: p<0.05, **: p<0.005 and ***: p<0.0005).

We next investigated the role of Cdk1 phosphorylation of Sam68 in apoptosis regulation using the Annexin V-APC PI flow cytometry assay (Supplementary Figure S5). Compared to untransfected cells, transfection of Sam68 WT increased the percentage of cells undergoing apoptosis as expected (from 14.2±1.5% to 22.9±1.3%), but this increase was significantly smaller upon transfection of the phospho-mimetic mutants (16.5±2.5, 15.1±0.4, and 17.5±2.1 for T33E, T317E and T33E/T317E, respectively) (Figure 7C, D).

Next, we investigated the rate of proliferation of HCT-116 cells upon transfection of wild-type Sam68 or phospho-mimetic mutants by live cell imaging taking images every 2 hours for 24 hours and counting the number of GFP-positive cells. Consistent with a reduction in apoptosis regulation, the rate of cell proliferation was significantly higher with the phospho-mimetic mutants (24-hour normalized cell proliferation ratio of 0.84±0.03, 1.03±0.07, 0.86±0.09 for T33E, T317E, T33E/T317E, respectively) compared to the wild-type Sam68 (0.68±0.03) (Figure 7E, Supplementary Figure S6).

Altogether, our data demonstrate that phosphorylation of Sam68 at T33 and T317 alters its regulatory role in cell cycle progression and apoptosis leading to increased proliferation of HCT-116 cells.

## DISCUSSION

It is well established that regulation of alternative splicing by splicing factors is modulated by cell signaling pathways (2,3). Indeed the large majority of splicing factors possess intrinsically disordered regions (IDRs) that are target sites for PTMs, and these modifications affect the function of splicing factors through various mechanisms, such as changes in cellular localization or RNA binding affinity (47,48). However in most cases, the molecular details of how PTMs affect splicing factor functions remain unclear. In the case of Sam68, it is known that phosphorylation, acetylation and methylation regulates its localization and function in alternative splicing (reviewed in (26)), but the effects of these modifications at the atomic level are largely unknown. Here we have investigated the phosphorylation of Sam68 by Cdk1/Cyclin B at the atomic level and demonstrated that Cdk1 phosphorylates the T33 and T317 residues located in the N-terminal and C-terminal IDRs of Sam68, respectively. We further show that phosphorylation of these residues reduces the binding of these regions to RNA and, most likely as a consequence, alters Sam68 localization and reduces its splicing activity.

How the phosphorylation of Sam68 T33 and T317 reduces the RNA binding affinity of the N-terminal and C-terminal regions of Sam68 remains to be elucidated. One possibility is that T33 and T317 are directly involved in RNA binding and that the addition of a negatively-charged phosphate group induces an electrostatic repulsion with the negatively-charged RNA. Another possibility is that Sam68 N-terminal and C-terminal regions bind RNA through their arginine side-chains and, upon phosphorylation, the phosphate group forms a salt bridge with nearby arginine residues, preventing them from binding to RNA. Indeed it has been shown that phosphorylated amino acids can form very stable salt bridges with two arginine guanidino groups (49). This would be consistent with our NMR data showing that both RNA and phosphorylation induces chemical shift changes of the arginine side chain Nε (supplementary Figure S7) and that both T33 and T317 are surrounded by nearby arginines, R31 and R36 in the N-terminal region and R315 and R320 in the C-terminal region.

Together, our data suggest a mechanistic model for the role of Cdk1 phosphorylation in Sam68 functions (Figure 8). In normal interphase cells, T33 and T317 of Sam68 are unphosphorylated allowing the N-terminal and C-terminal regions of Sam68 to bind RNA, most probably through their RG-rich regions. In this state, the STAR domain of Sam68 binds its pre-mRNA substrate specifically by recognizing the sequence (A/U)AA-N>15-(A/U)AA with a weak to medium affinity (dissociation constant in the micromolar range) (18). The primary binding of the STAR domain to the pre-mRNA brings the RG-rich regions of Sam68 N-terminal and C-terminal IDRs in close proximity to the RNA. As the STAR domain also promotes dimerization, two N-terminal and two C-terminal IDRs can bind the RNA, anchoring the protein to its target pre-mRNA (Figure 8, top), leading to high affinity RNA binding (dissociation constant in the low nanomolar range as reported previously for full-length Sam68 (41,42)). Upon mitotic entry - or potentially in certain cancers - Cdk1 activity increases, leading to Sam68 phosphorylation at T33 and T317. These phosphorylation events reduce the RNA binding affinity and induce the release of Sam68 N-terminal and C-terminal IDRs from the pre-mRNA (Figure 8, middle). The weakened affinity of phosphorylated Sam68 for the RNA reduces its ability to compete with other splicing factors and lead to its release from the pre-mRNA (Figure 8, bottom) and consequently its cellular relocalisation. In a normal cell, this could be important to prevent deleterious splicing events in mitosis. However, in a cancer cell, this could lead to reduced apoptosis and increased proliferation.

**Figure 8:**
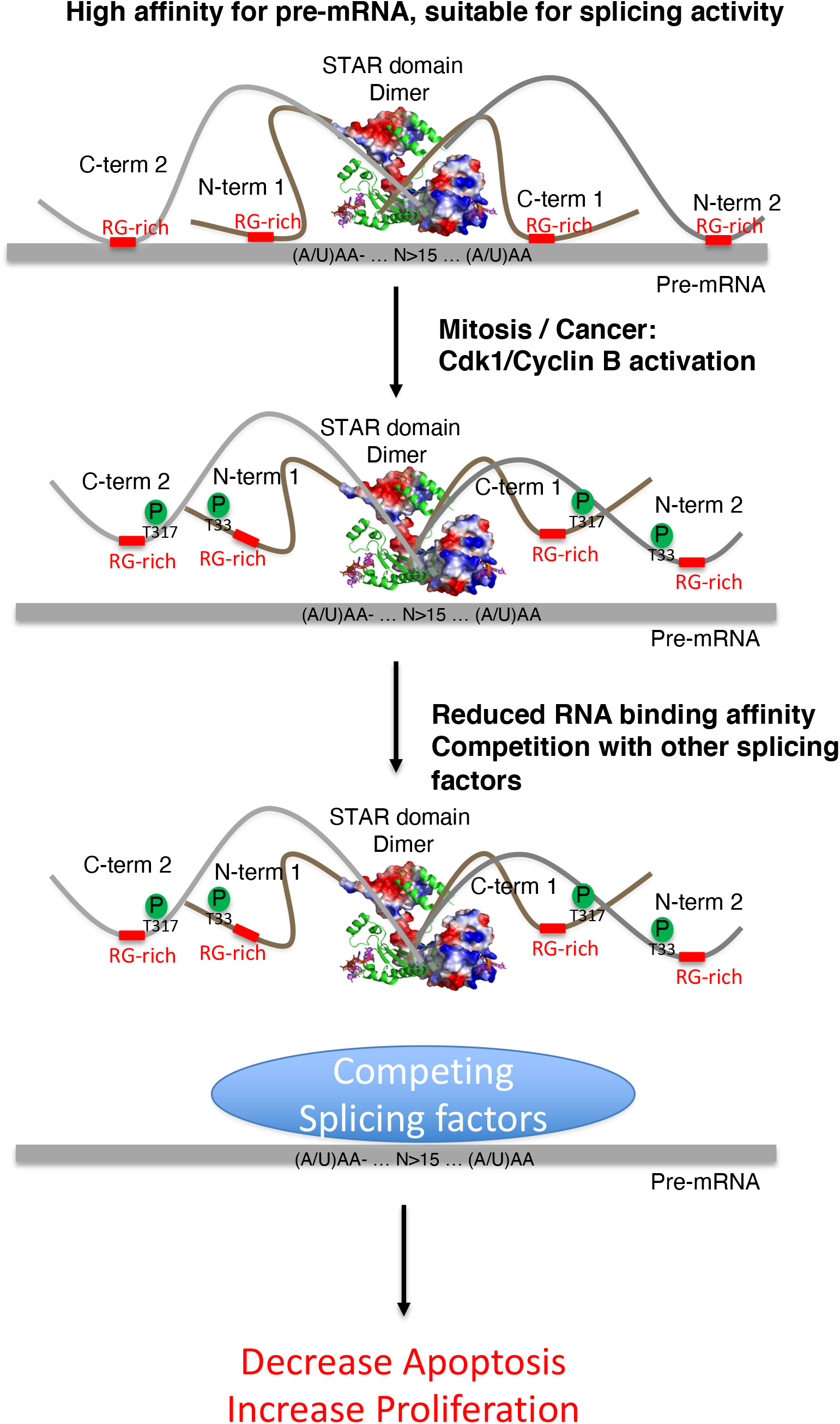
Model of splicing regulation by Cdk-1 mediated phosphorylation of Sam68. In interphase, Cdk-1 is inactive, so T33 and T317 of Sam68 are unphosphorylated. Sam68 would therefore bind RNA specifically through its QUA1-KH homodimerization domain with dissociation constant (Kd) in the low micromolar range and the N-terminal and C-terminal regions would anchor the protein to the RNA, increasing the affinity of full-length Sam68 to RNA (Kd in the low nanomolar range). During mitosis or in cancer cells, activation of Cdk-1/cyclin B induces the phosphorylation of T33 and T317, leading to the dissociation of the N-term and C-term anchoring region from the RNA and therefore, a weakening in RNA binding by full-length Sam68 making Sam68 less effective at competing with other splicing factors for RNA binding. As a consequence, Sam68 activity in splicing regulation is reduced, leading to a decrease in cell apoptosis and an increase in proliferation.

## MATERIAL AND METHODS

### Protein and RNA constructs

Sam68 constructs were cloned by the University of Leicester cloning service (Protex, https://le.ac.uk/mcb/facilities-and-technologies/protex). Full-length His-tag/Flag-tag Sam68 was cloned into the pLEICS-12 vector, Full-length GFP-Sam68 into the pLEICS-25 vector, His-Tag Sam68 N-terminal region (aa 1-96) into the pLEICS-01 vector and GST-Sam68 C-terminal region (267-368) into pGEX-6P-2 (https://www.genscript.com/express-cloning-vector-list.html). Sam68 C-terminal cDNA was synthetized with optimized codon composition for bacterial expression (Genscript).

RNA oligonucleotides were purchased from Dharmacon, Horizon Discovery.

Bclx and CD44-v5 minigenes were cloned in EGFP-C1 plasmid following the CMV promoter using ExoIII (NEB) enzyme following manufacturer instructions. CD44 minigene was cloned in two steps. Firstly, a DNA sequence comprising β-globin exon 2, intron 2 and exon 3 was cloned in AgeI and HindIII restriction sites of EGFP-C1 plasmid using primers Bg-AgeI-Fwd (agatccgctagcgctACCGGTGGGCTGCTGGTTGTCTACCCATGG) and Bg-HindIII-Rev (aattcgaacttgagcGAATTCAACTTACCTGCCAAAATGATGAGAC). Secondly, CD44 variable exon 5 (v5, 117 bp long) with flanking intron sequences (238 bp upstream and 270 bp downstream) was PCR amplified from genomic DNA using primers CD44-Fwd (AAATTCATGTTATATGGTCGACAGCCAACAGCCCTACAAATGTTAG) and CD44-Rev (AACATGGTTAGCAGAGTCGACACCCTTAGGAACCATTAACAC), and cloned into the SalI restriction site positioned in the β-globin intron 2. Bcl-x minigene(50) was PCR amplified using primers Bcl-X-Fwd (TAGTGAACCGTCAGATCCGCTAGCGCTACCGGTGGGAGGTGATCCCCATGGCAG) and Bcl-X-Rev (GTCGACTGCAGAATTCGAAGCTTACTTACCTGGCCACAGTCAT) and cloned in AgeI and HindIII sites of the EGFP-C1 plasmid using ExoIII enzyme.

### Bacterial expression of Sam68 N-terminal and C-terminal regions

The expression vectors (pLEICS-01 and pGEX-6P-2) were transformed into *Escherichia coli* Rosetta(DE3) cells. Cells were grown at 37*°*C to an absorbance of 0.8 and incubated with 0.5 mM IPTG at 20*°*C overnight. His-Tag Sam68 N-terminal region was purified using a nickel affinity chromatography column. The His-tag was removed by incubating the protein with TEV protease in the presence of 0.5 mM EDTA and 1mM DTT at 15*°*C overnight. The protein sample was buffer exchanged using PD-10 column (G25 resin, cutoff of 7 kDa; GE Healtcare) against 50 mM sodium phosphate buffer pH 7.6, 150 mM NaCl and then loaded on a nickel affinity chromatography column. The cleaved protein was then recovered in the flow-through. GST-Sam68 C-terminal region was first purified using glutathione affinity column. The eluted protein was incubated with PreScission protease at 4*°*C overnight. The cleaved protein was then purified by a Superdex 75 size exclusion column using 50 mM sodium phosphate buffer pH 7.0, 150 mM NaCl as running buffer.

All the purification steps were performed in the presence of complete protease inhibitor cocktail (Roche). Uniformly ^15^N, ^13^C labelled proteins were expressed in M9 medium containing 1g of ^15^NH_4_Cl and 2g of ^13^C-glucose per litre of culture as the sole source of carbon and nitrogen.

### Site-directed mutagenesis

Site-directed mutagenesis was carried out using extension PCR with back-to-back oriented primers, with the site of mutation positioned at the beginning of the forward primer. PCR reaction was carried out and the linear product of the PCR reaction was purified, ligated and transformed into DH5α. Colonies were screened and verified by Sanger sequencing.

### Cell culture

HEK293T and HCT116 were grown in Dulbecco’s Modified Eagle Medium (DMEM) supplemented with GlutaMAX™ (Gibco), 10 % fetal bovine serum (Gibco), 1% v/v penicillin and streptomycin (Gibco) in 5% CO_2_ incubator.

### Transfection

HCT116 cells were transfected using JetPrime(Polyplus) and HEK293 using Fugene (Promega) using a standard protocol. Cells were passaged 24 hours before transfection. 1.5-3μg of DNA was used to transfect 0.2×10^6^ passaged cells. Transfection efficiency was assessed by visualization of the GFP signal, and all experiments were done using cell cultures displaying >70% GFP positive cells (Supplementary Figure S1).

### Mass Spectrometry

Cells were passaged 24 hours before transfection with Flag-Sam68. For unsynchronized samples, cells were harvested 48h after transfection. For mass spectrometry analysis at different stages of the cell cycle, cells were pre-synchronized with 2mM thymidine 8 hours after transfection, and were released from the arrest by three washes with PBS followed by addition of fresh media 16 hours after thymine addition. For mitotic samples, cells were synchronized by addition of 0.5μg/ml nocodazole 8 hours after release, and harvested 16 hours after nocodazole addition. For G1 samples, cells were synchronized by addition of 1μg/ml nocodazole 8 hours after release from thymidine and then washed three times with PBS followed by addition of fresh media and incubation for 6 hours prior to harvesting. For S phase samples, cells were synchronized by addition of 1mM hydroxyurea 8 hours after release from thymidine and then washed three times with PBS followed by addition of fresh media and incubation for 3 hours prior harvesting. Cells were lysed with RIPA buffer and FLAG-Sam68 was pulldown using M2 FLAG beads. Purified FLAG-Sam68 was separated on Tris-Glycine gels and digested by trypsin: Gel plugs were washed with B solution (200mM TEAB, 50% acetonitrile) three times, each time incubating for 20 minutes, then washed with 100% acetonitrile and incubated for 10 minutes. After acetonitrile removal, gel plugs were air dried for 10 minutes, following by incubation in DTT solution (10mM DTT, 50mM TEAB) for 30 minutes at 60°C. Then gel plugs were incubated in iodocetamide solution (100mM iodecetamide, 50mM TEAB), followed by two washes with B solution, each time incubating for 20 minutes, then washed once with 100% acetonitrile and incubated for 10 minutes. After acetonitrile removal, gel plugs were air dried for 10 minutes, incubated in trypsin solution (50ng trypsin, 50mM TEAB) overnight at 37°C. After incubation the supernatant was collected from the gel plugs into new tube (A). After addition of 25% Acetonitrile, 5% formic acid solution to gel plugs, samples were sonicated in water bath for 10 minutes followed by liquid collected into tube A. This step was repeated three times. Samples were concentrated using a speed vacuum centrifuge, resuspended in 2% acetonitrile, 0.1% formic acid. Reversed phase chromatography was used to separate tryptic peptides prior to mass spectrometric analysis. Two columns were utilised, an Acclaim PepMap μ-precolumn cartridge 300 μm i.d. x 5 mm 5 μm 100 Å and an Acclaim PepMap RSLC 75 μm x 50 cm 2 μm 100 Å (Thermo Scientific). The columns were installed on an Ultimate 3000 RSLCnano system (Dionex). Mobile phase buffer A was composed of 0.1% formic acid in water and mobile phase B 0.1 % formic acid in acetonitrile. Samples were loaded onto the μ-precolumn equilibrated in 2% aqueous acetonitrile containing 0.1% trifluoroacetic acid for 5 min at 10 μL min^-1^ after which peptides were eluted onto the analytical column at 250 nL min^-1^ by increasing the mobile phase B concentration from 4% B to 25% over 37 min, then to 35% B over 10 min, and to 90% B over 3 min, followed by a 10 min re-equilibration at 4% B. Eluting peptides were converted to gas-phase ions by means of electrospray ionization and analysed on a Thermo Orbitrap Fusion (Q-OT-qIT, Thermo Scientific). Survey scans of peptide precursors from 375 to 1575 m/z were performed at 120K resolution (at 200 m/z) with a 2 × 10^5^ ion count target. Tandem MS was performed by isolation at 1.2 Th using the quadrupole, HCD fragmentation with normalized collision energy of 33, and rapid scan MS analysis in the ion trap. The MS^2^ ion count target was set to 5×10^3^ and the max injection time was 200 ms. Precursors with charge state 2–6 were selected and sampled for MS^2^. The dynamic exclusion duration was set to 25 s with a 10 ppm tolerance around the selected precursor and its isotopes. Monoisotopic precursor selection was turned on. The instrument was run in top speed mode with 2 s cycles. Data were searched using Mascot (version 2.6.1, Matrix Science Ltd., UK) against the Homo sapiens reference proteome database (www.uniprot.org/proteomes), and results imported into Scaffold (version 5.1.0, Proteome Software Inc.). Spectra of modified peptides with Mascot delta score > 0 were manually inspected to confirm sites of modification.

### NMR

NMR experiments were recorded on a Bruker 600-MHz spectrometer equipped with triple-resonance 1H/13C/15N cryogenic probe. NMR measurements were performed in 50M sodium phosphate buffer, pH 7.0, 150 mM NaCl and 10% D_2_O. All NMR spectra were processed using TopSpin software and analysed by CcpNmr Analysis (51).

Sam68 N-term and C-term regions backbone assignments were done using the following triple resonance experiments: HNCO, CBCA(CO)MH and HNCACB. In order to get optimal signal with less peak overlaps, experiments were measured at 4°C.

(^1^H-^15^N)-HSQC spectra were recorded to probe the interaction between ^15^N labelled N-term or C-term regions and G8-5 unlabelled RNA. The weighted chemical shift changes, Δδ(^1^H,^15^N), were calculated using the following equation: Δδ(H,N) = (Δδ^2^_H_ + Δδ^2^_N_ × 0.159)^1/2^.

Dissociation constants were obtained by fitting the chemical shift perturbation data to the following equation: 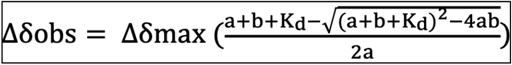, where Δδobs is the average weight of the chemical shifts in the free and bound states and Δδmax is the maximal signal change upon saturation. Kd is the dissociation constant, a and b are the total RNA and Sam68 C-term or N-term regions concentrations, respectively. Dissociation constants were calculated based on chemical shift perturbations of eight distinct backbone resonances or based on the arginine side chains resonances.

### Phosphorylation of Sam68 C-terminal and N-terminal regions

Phosphorylation of Sam68 N-term and C-term were obtained by incubation with CDK1/Cyclin B (Merck, catalogue 14-450) at a Cdk1:Sam68 ratio of 5:1000 at 20 °C, in the presence of 2 mM ATP, 5 mM MgCl2 and protease inhibitors in 50 mM sodium phosphate buffer pH 7.0 and 150 mM NaCl. First an initial (^1^H-^15^N)-HSQC was recorded at 4°C. Phosphorylation was then quantitatively monitored with consecutive 2D (^1^H-^15^N)-SOFAST-HMQC NMR experiments(52) for 16 hours at 20°C, and a final (^1^H-^15^N)-HSQC at 4°C. The time course of the normalized amplitude of phosphorylated T33 and T317 is sigmoidal rather than hyperbolic and was fitted using a Hill function.

### Fluorescence microscopy

HCT116 and HEK293 cells were seeded on 1.5mm coverslips (VWR) 24 hours prior to transfection with pLEICS25 wild-type or mutant GFP-Sam68. Transfected cells were cultured for 48 hours, washed with PBS and fixed with 4% paraformaldehyde for 10 minutes. Paraformaldehyde was removed by three washes with PBS (Life technologies), then DNA was stained with 300nM DAPI (ThermoScientific) for 5 minutes. Excess of DAPI was removed by washing cells three times with PBS. Coverslips were mounted on slides using 3% (w/v) n-propyl gallate (Sigma) in an 80% glycerol solution and sealed with clear nail varnish. Cells were imaged using Zeiss LSM 980 Airyscan 2 microscope in confocal mode with Plan-Apochromat 63x/1.40 Oil DIC f/ELYRA lens. 405nm laser was used to excite DAPI and was detected within 415-475nm, 488nm laser was used to excite GFP and was detected within 495-555nm. Sam68 localization patterns were assigned and counted manually for 50-100 cells per sample. Statistical analysis was done using GraphPad/Prism

### Splicing assays

The splicing activity of the Sam68 WT and mutant proteins were assessed in HCT116 cells using a minigene assay as previously described (53). In brief, HCT116 cells were transfected with WT or mutant GFP-Sam68 and *CD44 or Bcl-x* minigenes. Transfected cells were harvested after 48 hours. RNA was isolated and purified using Monarch Total RNA Miniprep Kit (Biolabs) following the manufacturer’s protocol. cDNA was transcribed using High-Capacity cDNA Reverse Transcription Kit (TermoFisher Scientific). BCL-X minigene was amplified with the forward primer 5′-AGTTTGAACTGCGGTACCGGCG-3′ and reverse primer 5′-TCATTTGTATTTCCAAGGAGTTAACCTC-3′. CD44 minigene was amplified with the forward primer 5′-CTGTCCTCTGCAAATGCTGTTTATGAAC-3′ and reverse primer 5′-AATAACCAGCACGTTGCCCAGGAG-3′. PCR products were separated on 1.5% agarose gel and visualized with a U:GENIUS gel imaging system (Syngene). Band intensities were quantified with ImageJ and statistical analysis was done using GraphPad/Prism.

### Cell cycle progression assays

Cell-cycle profiles were determined by measuring cellular DNA content by flow cytometry. 48 hours after transfection, cells were collected by trypsinization, washed with phosphate buffer saline (PBS) and fixed with ice cold 70% ethanol at -20°C overnight. Cells were washed three times with PBS and resuspended in 300μl PBS, then RNAse A (200 μg/ml final concentration) and propidium iodide (PI) (20 μg/ml final concentration) were added to the samples and incubated at 4°C overnight in the dark. Cell cycle profiles were determined using CytoFlex (Beckman) and analysed using FlowJo. Statistical analysis was done using GraphPad/Prism.

### Apoptosis assays

Apoptosis was determined using the Annexin V-APC PI kit (Biolegend). 48 hours after transfection, cells were collected by trypsinization and washed with phosphate buffered saline (PBS). The cells were resuspended in 300 μl of binding buffer, then 5μl of Annexin V-APC (stock concentration of 0.5 mg/ml) and 5μl propidium iodide (stock concentration of 4mg/ml) were added to the samples for 10 minutes and incubated in the dark. Apoptosis were determined using CytoFlex and analysed using FlowJo. Statistical analysis was done using GraphPad/Prism.

### Proliferation assays

24 hours before transfection, HCT116 cells were seeded as 1000 cells per well in 24-well plates. Cells in each well were transfected with 0.38μg of plasmid DNA. 48 hours after transfection, GFP signal was recorded by taking images of cells every 2 hours for 24 hours with LiveCite 2. GFP positive cells were counted manually at timepoints 48 and 72 hours after transfection, or counted at each time point using Trainable Weka Segmentation (54), ITCN plugin and macro in ImageJ (55). Statistical analysis was done using GraphPad/Prism.

## ACKNOWLEDGEMENT

We would like to acknowledge the support of Dr. F. Muskett and the Leicester NMR facility, Dr. K. Straatman and the Advanced Imaging Facility (RRID:SCR_020967). We also would like to thank I. Eperon, L. O’Regan, G. Thomas, and V. Smith for useful discussions.

## ACCESSION NUMBERS

NMR Chemical shift assignments of Sam68 Nter and Cter have been deposited to the BioMagResBank (BMRB) under accession numbers 51359 and 51360, respectively.

Mass spectrometry data have been deposited to PRIDE under accession numbers …

Flow cytometry data have been deposited to the FlowRepository under accession numbers FR-FCM-Z55E (associated with Supplementary Figure S2), FR-FCM-Z55H (associated with Supplementary Figure S4), and FR-FCM-Z55Z (associated with Supplementary Figure S5).

Mass spectrometry proteomics data have been deposited to the ProteomeXchange Consortium via the PRIDE (56) partner repository with the data set identifier PXD032716 and 10.6019/PXD032716.

## FIGURES LEGENDS

**Supplementary Figure S1:**
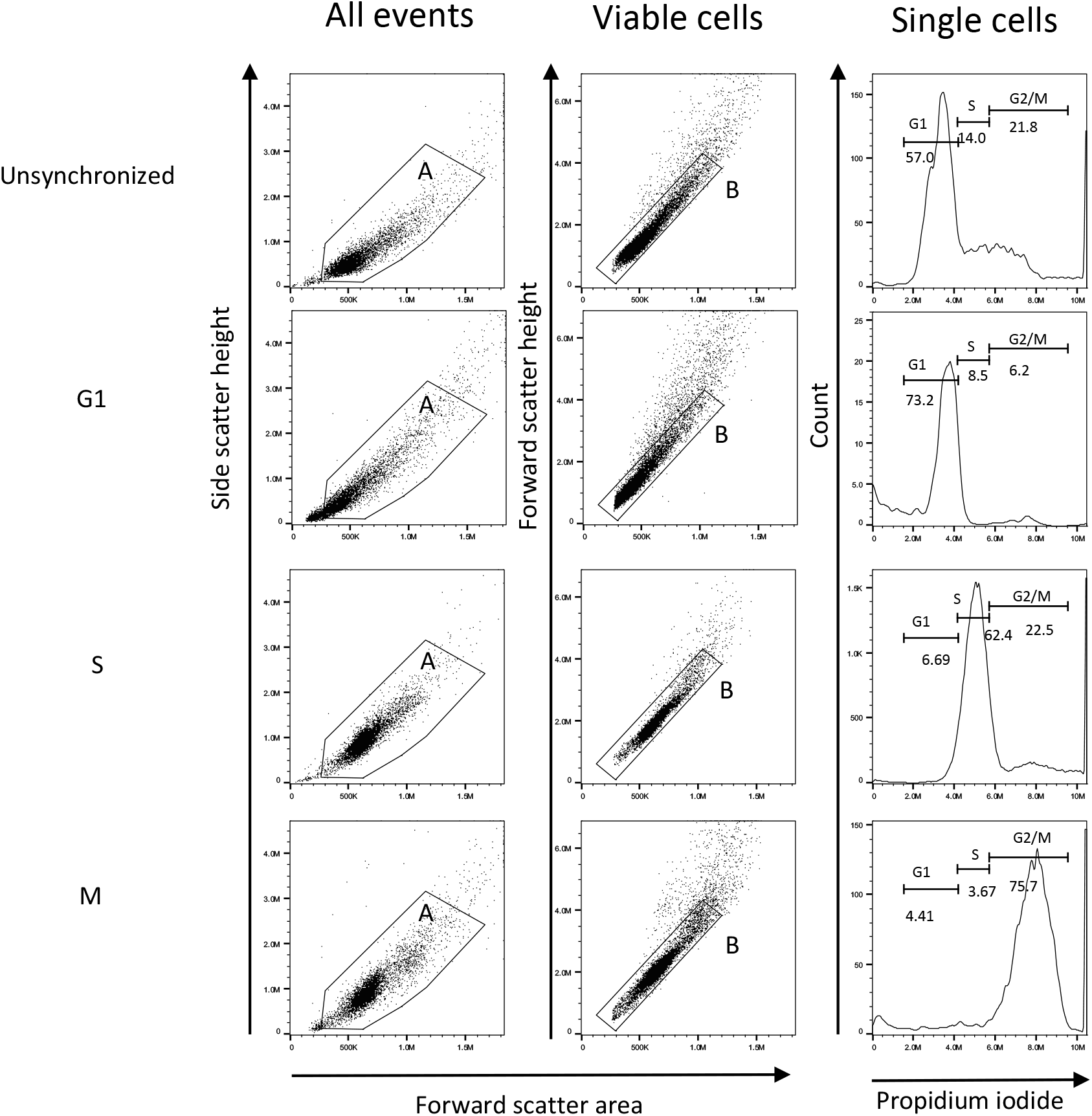
Cell cycle distribution in unsynchronized and synchronized HEK293 cells. All collected events were gated for viable cells (gate A) and single cells (gate B). Viable and single cells are gated from G1, S and G2/M phase

**Supplementary Figure S2:**
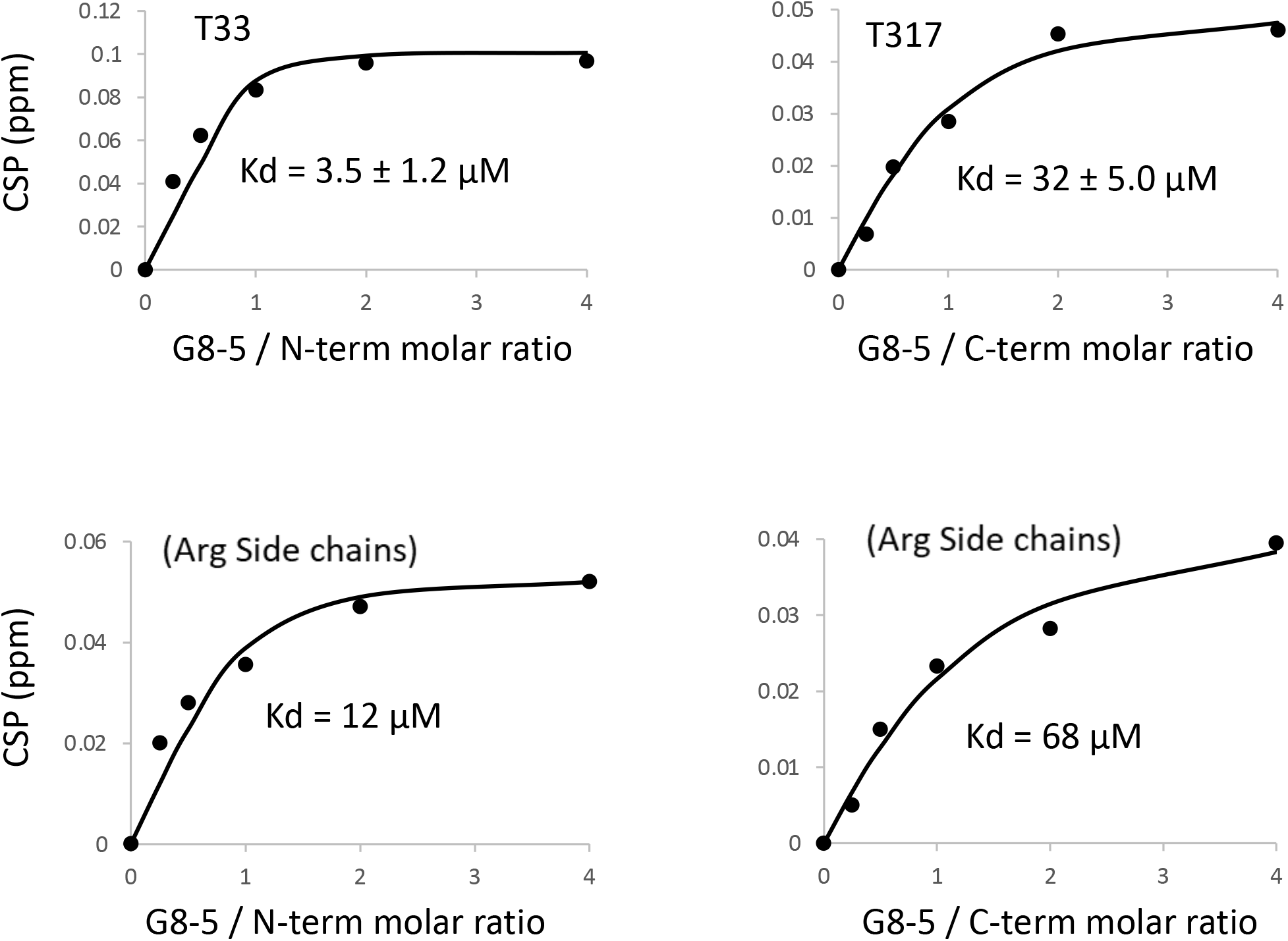
Estimation of Sam68 N-term (left) and C-term (right) dissociation constants upon interaction with the G8.5 RNA. For backbone atoms, CSP as a function of RNA/protein ratio are shown for T33 and T317 as examples. The Kd was estimated based on the CSP of eight backbone resonances. The CSP as a function of RNA/protein ratio is also displayed and the Kd estimated for arginine side chain resonances.

**Supplementary Figure S3:**
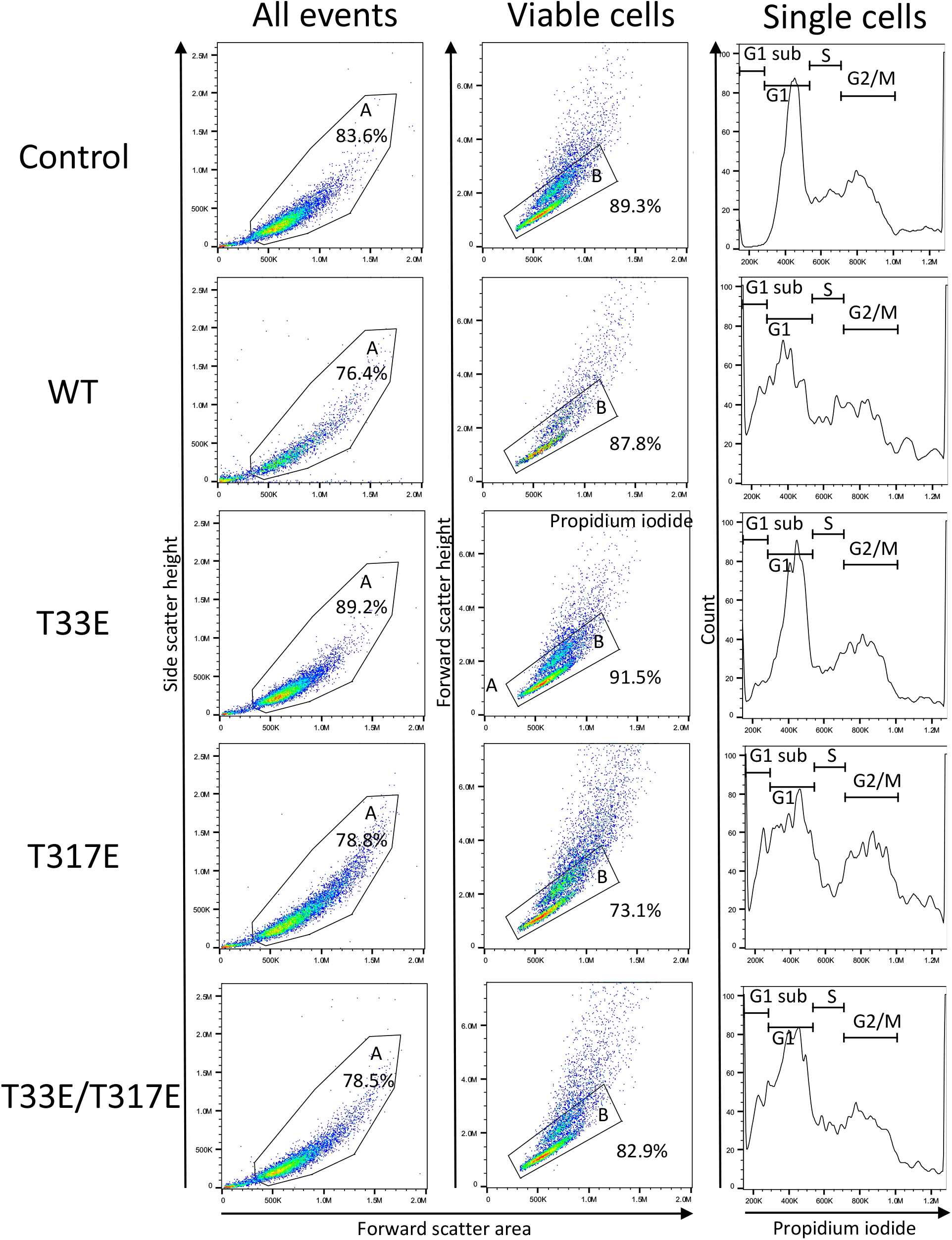
Cell cycle analysis by flow cytometry. All collected events were gated for viable cells (gate A) and single cells (gate B). Viable and single cells are gated for G1 sub, G1, S and G2/M phase.

**Supplementary Figure S4:**
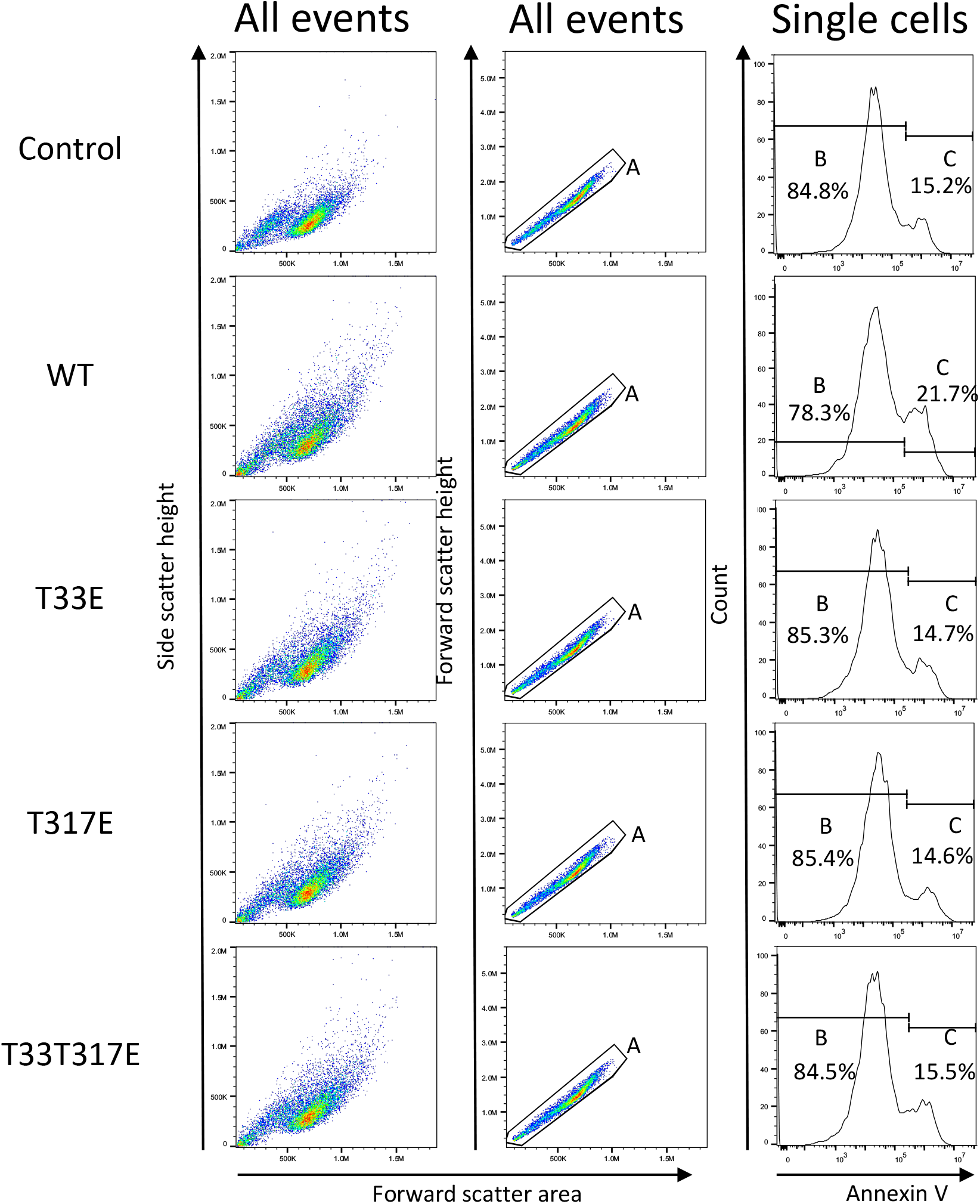
Apoptosis analysis by flow cytometry. All collected events were gated for single cells (gate A). Single cells are gated for viable (gate B) and apoptotic (gate C) cells.

**Supplementary Figure S5:**
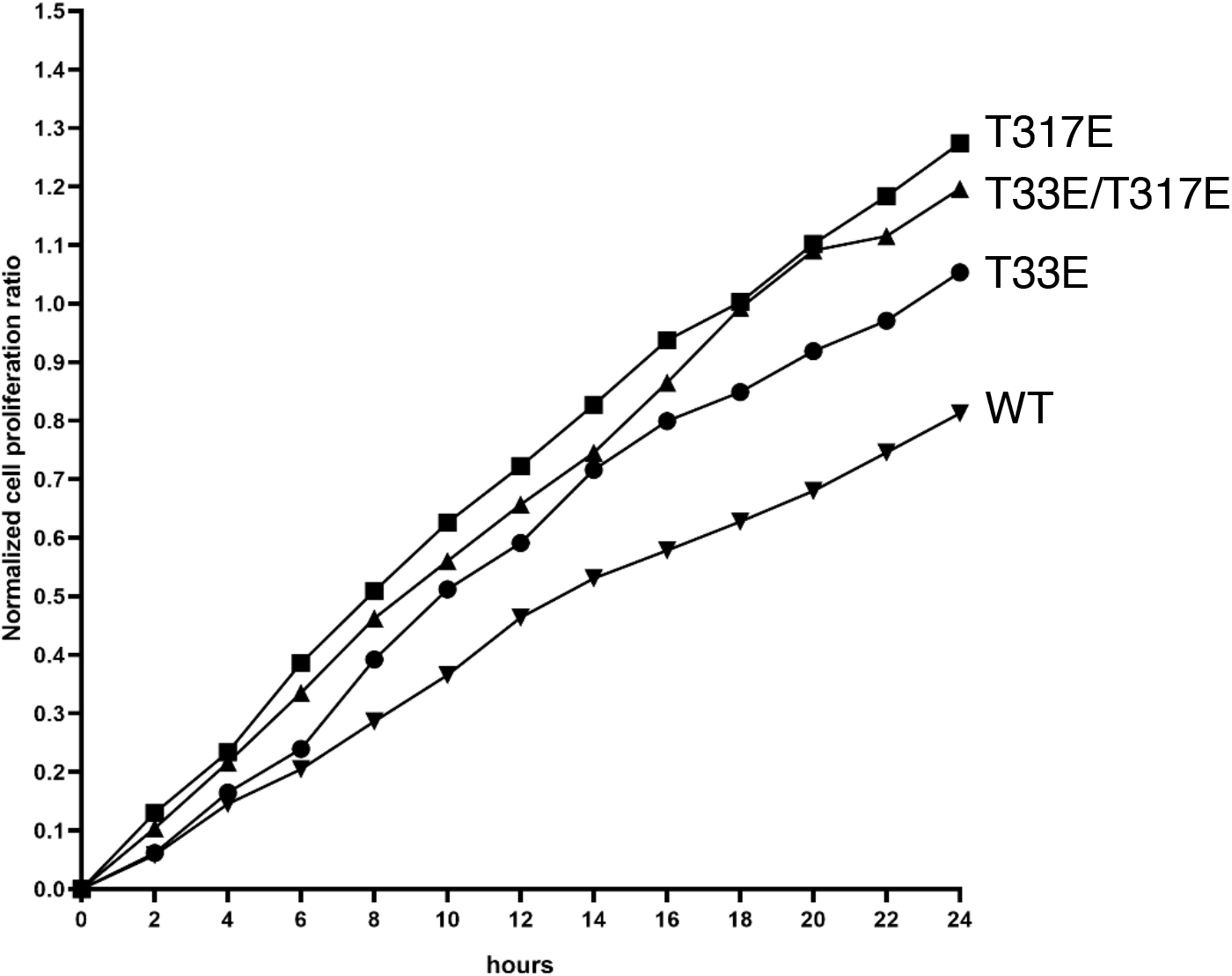
Proliferation curves following transfection with Sam68 WT and phospho-mimetic mutants. 48 hours after transfection of HCT116 cells with Sam68 WT or mutants, the number of GFP(+) cells were counted every 2 hours for 24 hours with LiveCite 2 using Trainable Weka Segmentation^52^, ITCN plugin and macro in ImageJ^53^.

**Supplementary Figure S6:**
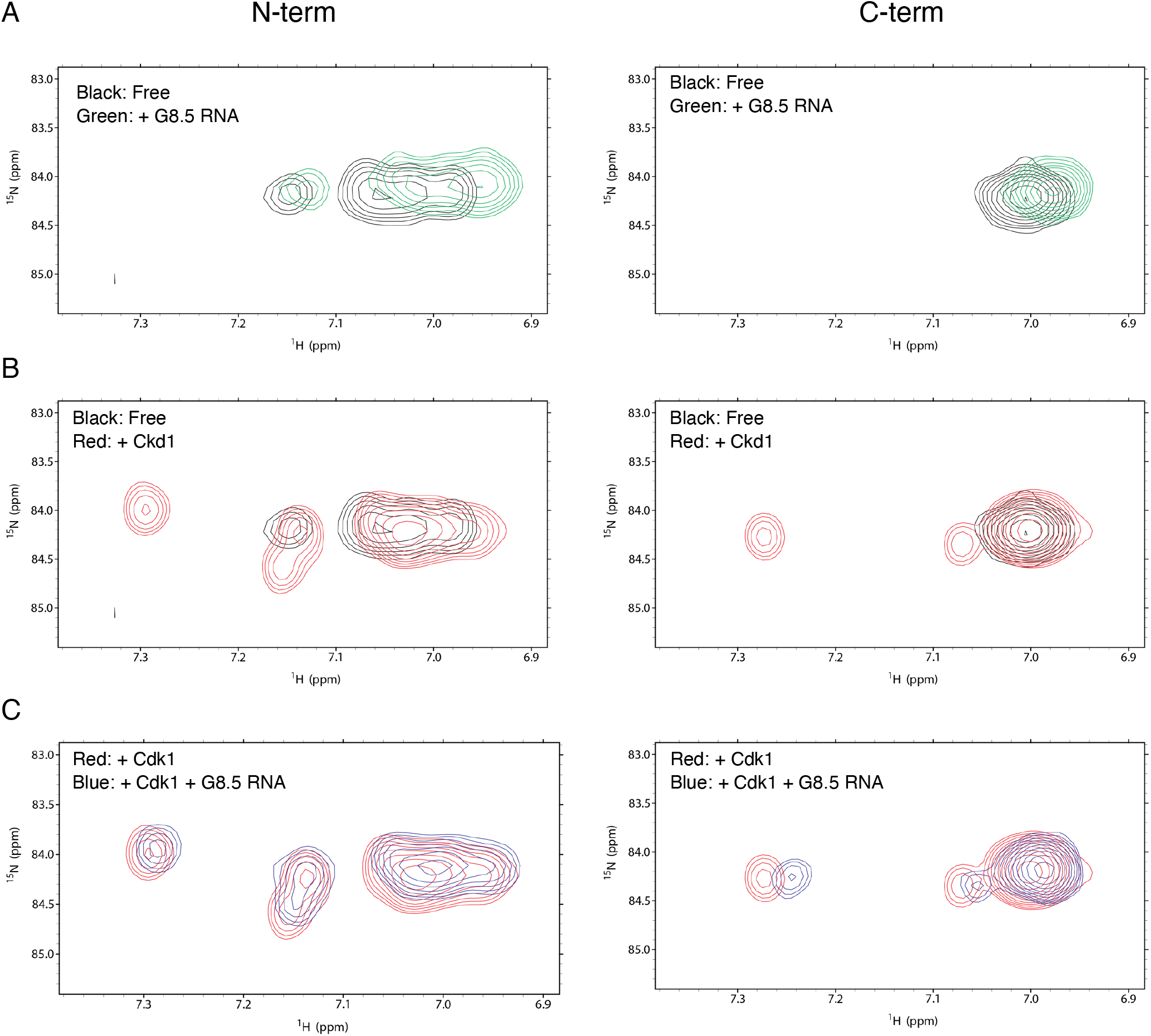
NMR resonances of arginine side chains upon RNA binding or Cdk1 phosphorylation. Overlay spectra of N-term (left and C-term (right) regions of Sam68. A) resonances in the absence (black) or presence (green) of 2 molar equivalents of G8.5 RNA; B) resonances before (black) and after 8 hours incubation with Cdk1, MgCl_2_ and ATP (red); C) resonances after Cdk1 phosphorylation in the absence (red) and presence (blue) of 2 molar equivalents of G8.5 RNA.

**Supplementary Figure S7:**
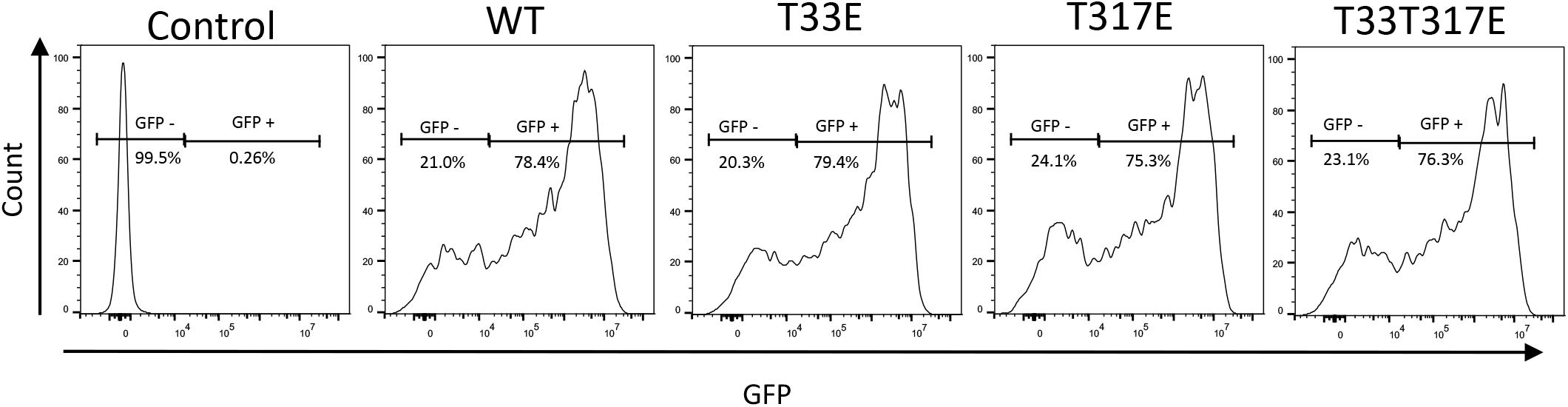
Analysis of transfection efficiency by flow cytometry. 48 hours after transfection of HCT116 cells with Sam68 WT or mutants, the number of GFP(+) nad GFP(-) cells were counted. Cells were used for additional experiments only if more than 75% of cells were GFP positive.

**Supplementary Figure S8:**
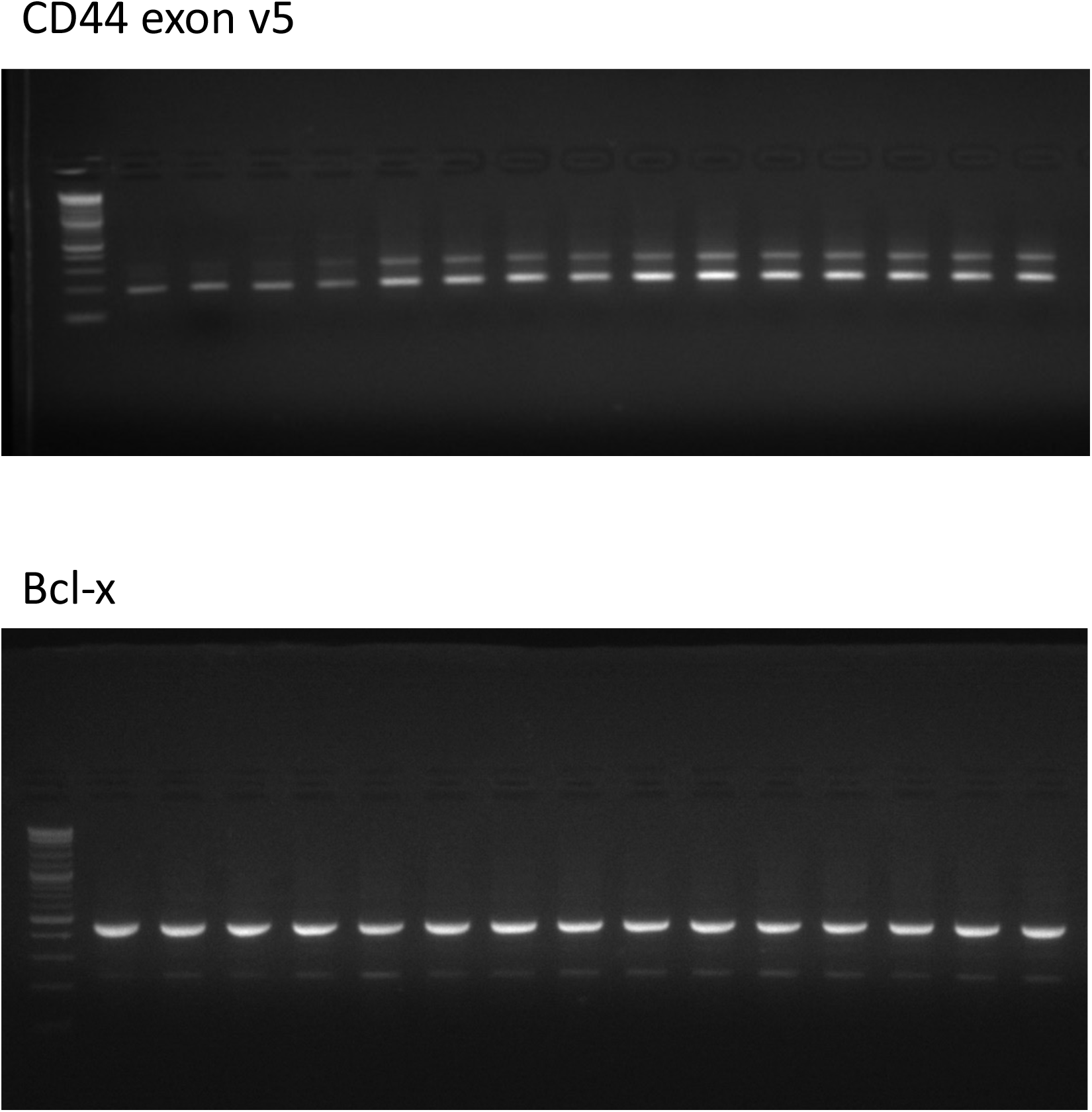
uncropped gels of splicing assays.

